# Rapid Cell Type-Specific Nascent Proteome Labeling in *Drosophila*

**DOI:** 10.1101/2022.10.03.510650

**Authors:** Stefanny Villalobos-Cantor, Ruth M. Barrett, Alec F. Condon, Alicia Arreola-Bustos, Kelsie M. Rodriguez, Michael S. Cohen, Ian Martin

## Abstract

Controlled protein synthesis is required to regulate gene expression and is often carried out in a cell type-specific manner. Protein synthesis is commonly measured by labeling the nascent proteome with amino acid analogs or isotope-containing amino acids. These methods have been difficult to implement *in vivo* as they require lengthy amino acid replacement procedures. O-propargyl-puromycin (OPP) is a puromycin analog that incorporates into nascent polypeptide chains. Through its terminal alkyne, OPP can be conjugated to a fluorophore-azide for directly visualizing nascent protein synthesis, or to a biotin-azide for capture and identification of newly-synthesized proteins. To achieve cell type-specific OPP incorporation, we developed phenylacetyl-OPP (PhAc-OPP), a puromycin analog harboring an enzyme-labile blocking group that can be removed by Penicillin G acylase (PGA). Here, we show that cell type-specific PGA expression in *Drosophila* can be used to achieve OPP labeling of newly-synthesized proteins in targeted cell populations within the brain. Following a brief 2-hour incubation of intact brains with PhAc-OPP, we observe robust imaging and affinity purification of OPP-labeled nascent proteins in PGA-targeted cell populations. We apply this method to show a pronounced age-related decline in neuronal protein synthesis in the fly brain, demonstrating the capability to quantitatively capture *in vivo* protein synthesis states using PhAc-OPP. This method, which we call POPPi (*P*GA-dependent *OPP i*ncorporation), should be applicable for rapidly visualizing protein synthesis and identifying nascent proteins synthesized under diverse physiological and pathological conditions with cellular specificity *in vivo*.

## INTRODUCTION

Controlled protein synthesis plays a fundamental role in orchestrating gene expression, and cellular protein amounts are thought to be primarily determined at the level of translation (1). Translational control is integral to cellular development and homeostasis and becomes dysregulated in numerous disease states including cancer (2), autism (3) and neurodegeneration (4). Within a complex organ like the brain, the specialized and distinct properties of neuronal and glial subtypes arise from variable expression of many protein components which are subject to translational control (5). Regulation of global protein synthesis as well as the modulated synthesis of specific proteins at the synapse is important in synaptic plasticity and memory formation (6).

Despite advances in assessing the transcriptome with cellular resolution, mRNA levels are an imprecise surrogate for protein abundance and methods to capture the nascent proteome of individual cell populations *in vivo* have lagged behind. Current methods employ indiscriminate protein synthesis labeling in all cells of a tissue followed by flow sorting of tagged cell suspensions from these tissues (7) or regional microdissection (8) and suffer from a lack of precision or substantial loss of protein. An alternative method for quantifying protein synthesis is through non-canonical amino acid (NCAA) protein labeling, using methionine analogs. This approach recently progressed toward cell-type specificity through expression of engineered methionyl-tRNA synthetase (MetRS) (9-11). Only mutant MetRS can charge methionyl-tRNA with azidonorleucine (ANL) which is amenable to protein capture by BONCAT (biorthogonal non-canonical amino acid tagging) and visualization by FUNCAT (fluorescent non-canonical amino acid tagging) (9-11). Hence, nascent protein labeling is restricted to cells that express mutant MetRS. However, this labeling strategy requires extended dietary methionine depletion and lengthy ANL feeding in mice or 1-2d of ANL feeding in flies and chronic feeding is associated with significant developmental toxicity and behavioral deficits (9, 10).

A potentially more efficient approach to protein synthesis labeling is through the use of puromycin analogs (12, 13). Puromycin is structurally similar to tyrosyl tRNA, yet its incorporation is not amino acid-specific, hence in contrast to NCAA, its incorporation into nascent proteins is not biased by their sequence or extent of methionine content.

This facilitates a more uniform incorporation into newly-synthesized proteins (14), and therefore a more reliable assessment of global protein synthesis. Addition of an O-propargyl group to puromycin allows visualization or capture of newly-synthesized protein via click-chemistry conjugation to a fluorophore-azide or a biotin-tagged azide, respectively, and has been successfully demonstrated in cultured cells (13) and in cells isolated from OP-puromycin-injected animals (7). We developed an analog of OP-puromycin (OPP) called phenylacetyl-OP (PhAc-OPP) that harbors an enzyme-labile blocking group (12). This blocking group renders OPP incapable of nascent protein incorporation until its removal by the *E. coli* enzyme penicillin G acylase (PGA) (12).

Hence, targeted expression of PGA can, in principle, be used to limit OPP proteome labeling to individual cell populations within animal tissues. To directly test this, we generated PGA-transgenic *Drosophila* and developed a method that we call POPPi (PGA-dependent OPP incorporation) for rapid, cell type-specific labeling of protein synthesis in intact fly brains. Here, we show that POPPi is a versatile labeling strategy which can achieve efficient visualization and identification of newly-synthesized proteins in a cell population of interest. Our approach enables cell-specific nascent proteome labeling from complex brain tissue within just a few hours, making it a powerful and efficient tool for examining the role of translational control in different physiological and pathological states.

## RESULTS

We previously demonstrated that neuronal PGA expression successfully converts PhAc-OP-puromycin (PhAc-OPP) to OP-puromycin (OPP), thus enabling OPP labeling of nascent proteins in cultured primary mouse neurons ((12) and Figure 1-figure supplement 1A). Toward cell type-specific protein labeling in complex nervous system tissue *in vivo*, we generated a transgenic PGA fly line that can express FLAG-tagged PGA when crossed to any of the commonly available fly lines expressing a cell type-specific GAL4 driver of choice (Figure 1-figure supplement 1B). Following ubiquitous PGA expression, we assessed PGA levels in embryos and the brains of larvae, pupae and adults. PGA expression is detectable in both developing and adult flies, with highest expression levels seen in larval and pupal brains, followed by adult brains and embryos (Figure 1-figure supplement 1C). We reasoned that cell type-specific nascent protein labeling in the CNS might be efficiently achieved using intact *Drosophila* brain explants. *Drosophila* whole brains isolated and maintained *ex vivo* are remarkably stable, exhibiting sustained neuronal morphology and physiological properties for many hours (15-22). Fly brain explants have been used to study neuronal activity such as response to odor stimulation (19, 20), axon remodeling (16, 18), neuronal wiring (21), neural stem cell proliferation (22) and protein synthesis (17). To examine whether isolated adult fly brains exhibit stable protein synthesis, we generated fly brain preparations and assessed ^35^S-methionine/cysteine incorporation levels over eight hours. We observe no significant change in global protein synthesis rates over this time period (Figure 1A), suggesting that protein synthesis is stable in newly-isolated whole brain preparations and that these may therefore be suitable for capturing *in vivo* protein synthesis states.

**Figure 1.**
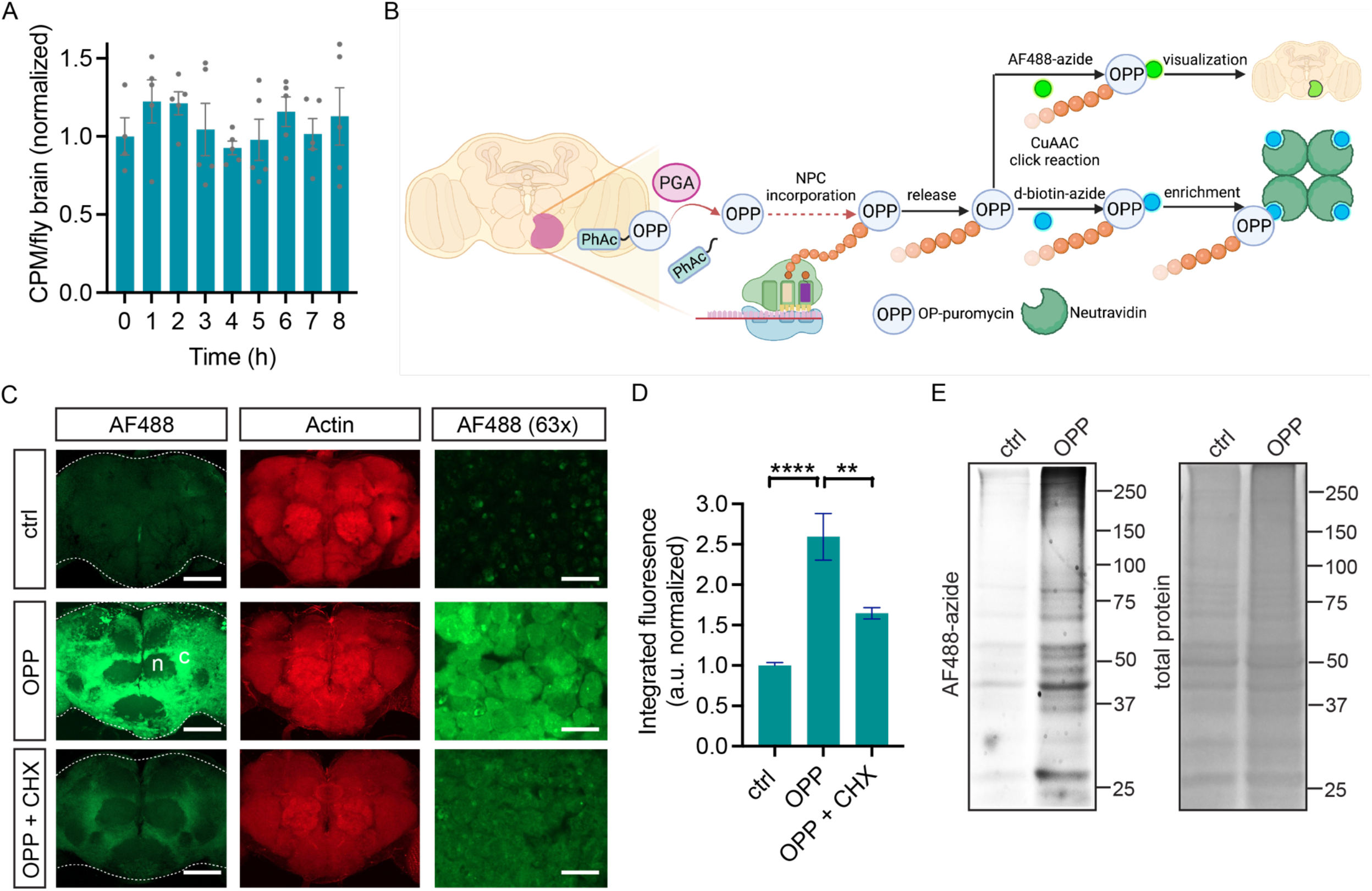
OPP labeling in *Drosophila* brain. A, global protein synthesis (^35^S-met/cys incorporation) is unaltered for 8h in newly-isolated *w*^*1118*^ whole brain preparations (ANOVA, n.s. n = 4-5 groups of 8-10 brains per timepoint). B, Schematic illustrating cell type-specific protein synthesis labeling by POPPi for visualization or capture of the nascent proteome. Spatially targeted PGA expression catalyzes PhAc-OPP blocking group removal, liberating OPP for incorporation into nascent polypeptide chains (NPC). OP-puromycilated proteins can be visualized by confocal microscopy following conjugation to a fluorescent-azide or enriched following conjugation to desthiobiotin-azide. C, newly-synthesized protein (from *w*^*1118*^ brains) visualized by AF488-azide after OPP incubation but not without OPP (ctrl) and diminished signal with cycloheximide (CHX). Labeling appears stronger in cell bodies within the cell cortex (c) than in the neuropil (n). Higher magnification (63x) images from the cell cortex. Scale bars are 60 μM (or 10 μM for 63x). D, quantitation of C, revealing a significant effect of treatment group (ANOVA, p<0.0001, Bonferroni post-test, ** p <0.01, **** p <0.0001, n =8-11 brains/group). E, in-gel fluorescence of brain protein extracts, ctrl is no OPP. Data are mean ± SEM. Schematic in B created on Biorender. See also Figure 1-figure supplements 1 and 2.

### *Drosophila* brain nascent proteomes can be tagged by OPP

Two major goals for cell-specific protein synthesis labeling are the visualization and identification of nascent proteomes in a cell type of interest. We hypothesized that this could be accomplished in *Drosophila* brain through OPP labeling of nascent polypeptide chains (NPC) followed by click-chemistry conjugation to either a fluorophore-azide for protein visualization or a biotin-azide for protein capture (schematic in Figure 1B). We first assessed whether unblocked OPP can label newly-synthesized proteins in *Drosophila* brain explants. Widespread OPP incorporation into newly-synthesized protein is clearly visible across the adult brain when coupled to Alexa Fluor 488-azide for detection (Figure 1C). The extent of incorporation is dependent on OPP concentration and slightly on incubation time (Figure 1-figure supplement 2) and is blocked by the protein synthesis inhibitor cycloheximide (Figures 1C and D), indicating that it is protein synthesis-dependent. Labeling is most prominently seen in the cell cortex, the location of cell bodies within the fly brain, consistent with the majority of protein synthesis occurring within the cell soma (Figure 1C). Theoretically, the protein synthesis imaged through this approach could represent OPP incorporation into a wide array of translating proteins or be restricted to a few highly-expressed proteins. In-gel fluorescence assessment of electrophoretically-separated brain extracts reveals that OPP-labeled proteins span the whole molecular weight range, while a minor degree of non-specific background fluorophore incorporation is seen in the absence of OPP (Figure 1E). This finding suggests that OPP is efficiently and unbiasedly incorporated into a broad set of proteins, which is also consistent with prior studies from OPP-labeling of cultured human cells (23).

### Rapid cell type-specific protein synthesis labeling with PhAc-OPP

We next assessed if targeted PGA expression can promote PhAc-OPP unblocking and OPP labeling of newly-synthesized proteins in cell populations of interest. We expressed PGA pan-neuronally in *Drosophila* brain using the *elavC155-GAL4* driver, which we confirmed by immunoblot (Figure 2-figure supplement 1) and then treated freshly-isolated brain explants with PhAc-OPP for 2h. Robust neuronal protein synthesis labeling is seen following PhAc-OPP treatment, but not in the absence of PhAc-OPP (Figure 2A) or in PhAc-OPP treated brains in the absence of GAL4-driven PGA expression (Figure 2-figure supplement 1B). The extent of labeling is time-dependent, although appears to be maximal at around 2h (Figure 2B). This is consistent with a prior study of OPP labeling in cells (13), and with a scenario that OPP-peptide conjugates eventually undergo turnover. When PGA expression was restricted to dopamine neurons using *TH-GAL4*, newly-synthesized proteins are clearly visible in the soma of dopamine neurons within intact brain explants briefly treated with PhAc-OPP, but not in the surrounding tissue (Figure 2C). PGA expression targeted to glia via *repo-GAL4*, also confirmed via immunoblot (Figure 2-figure supplement 1A), results in visible nascent protein labeling in glial cells (co-labeled with mCherry) upon brief PhAc-OPP treatment, but not in the absence of PhAc-OPP (Figure 2D). Hence, POPPi can efficiently label newly-synthesized proteins in neuronal and glial cell populations within the fly brain.

**Figure 2.**
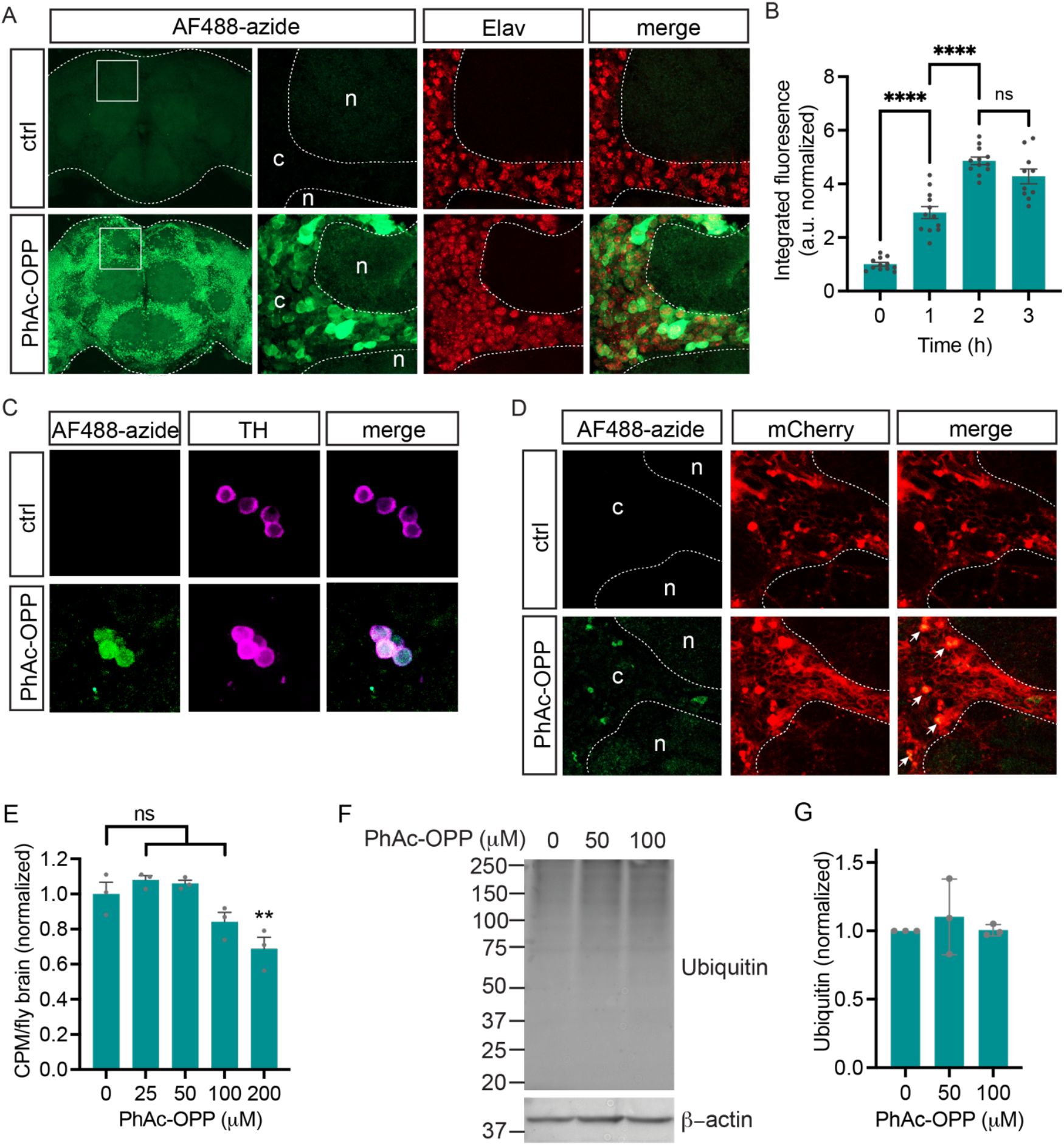
Cell type-specific protein synthesis labeling with PhAc-OPP. A, brains from *elavC155-GAL4;UAS-PGA* flies incubated with PhAc-OPP show widespread protein synthesis labeling in neurons, labeled with Elav neuronal nucleus marker. No labeling in vehicle-treated brains (ctrl). Labeling appears highest in the cell bodies of the cell cortex (c) and minimal in the neuropil (n). B, AF488-azide quantitation after varying durations of PhAc-OPP labeling in pan-neuronal PGA expressing flies. Significant effect of PhAc-OPP incubation time on labeling (ANOVA, Bonferroni post-test, **** p <0.0001, n =10-12 brains per group). C, protein synthesis labeling in isolated brains from flies expressing PGA in dopamine neurons (*TH-GAL4/UAS-PGA*). TH, tyrosine hydroxylase. D, protein synthesis labeling following pan-glial expression of PGA and a membrane-tethered mCherry reporter (*Repo-GAL4, UAS-mCD8::mCherry/UAS-PGA*). Glia cell bodies and neuron-encapsulating surface areas are mCherry-positive. Arrows indicate glia cell bodies positive for both mCherry and AF488, indicative of glial protein synthesis labeling. Controls in A, C and D are vehicle-treated brains. E, significant effect of 200 μM PhAc-OPP on global protein synthesis in flies expressing PGA ubiquitously via *Actin5C-GAL4* (ANOVA, Bonferroni post-test, ** p <0.01, n = 3 groups of 8-10 brains/group). F, no significant effect of PhAc-OPP on total protein ubiquitination in flies expressing PGA ubiquitously, quantified in G (ANOVA, n =3 groups of 20 brains/group). Data are mean ± SEM. See also Figure 2-figure supplement 1.

Since OPP incorporation into NPC results in peptide chain termination, we queried whether PhAc-OPP treatment diminishes overall protein synthesis rates. To assess this, we performed ^35^S-methionine labeling in parallel to PhAc-OPP treatment of brain explants expressing PGA ubiquitously. We found that PhAc-OPP causes a dose-dependent inhibition of global protein synthesis rates but that this inhibition was minimal up to 100 μM PhAc-OPP (Figure 2E) which is the standard concentration used in our labeling experiments. A significant disruption of protein synthesis is only seen above 100 μM PhAc-OPP (Figure 2E). Additionally, protein ubiquitination levels are not affected by incubating brains in up to 100 μM PhAc-OPP for 2 hours (Figures 2F and G), consistent with PhAc-OPP not having a major impact on protein turnover under these conditions.

### Rapid cell type-specific nascent proteome capture with PhAc-OPP

As observed upon treating brain explants with OPP (Figure 1E), we find that PGA-dependent unblocking of PhAc-OPP in all neurons or in all glia of adult fly brains gives rise to OPP-labeled proteins that span a wide molecular weight range consistent with broad incorporation into the nascent proteome (Figures 3A and C). In support of widespread incorporation into NPC, we were able to label specific neuronal proteins of interest when pan-neuronal PGA expression was coupled to PhAc-OPP treatment and brain lysates were clicked to desthiobiotin azide for affinity purification of the neuronal proteome (Figure 3B). Three crucial synaptic proteins involved in neurotransmitter release, namely Bruchpilot (ERC2 ortholog), Synapsin and Syntaxin, are all substantially enriched in the OPP-labeled neuronal proteome following affinity purification (Figure 3B). Similarly, glial PGA expression coupled to PhAc-OPP incubation and affinity purification with desthiobiotin azide yields strong enrichment of the glial-specific protein Draper (ortholog of the mammalian engulfment receptor MEGF10) (Figure 3D). Conversely, the neuronal SNARE complex protein Syntaxin is not enriched upon pan-glial PGA expression, supporting the conclusion that proteome labeling is restricted to the cell population expressing PGA.

**Figure 3.**
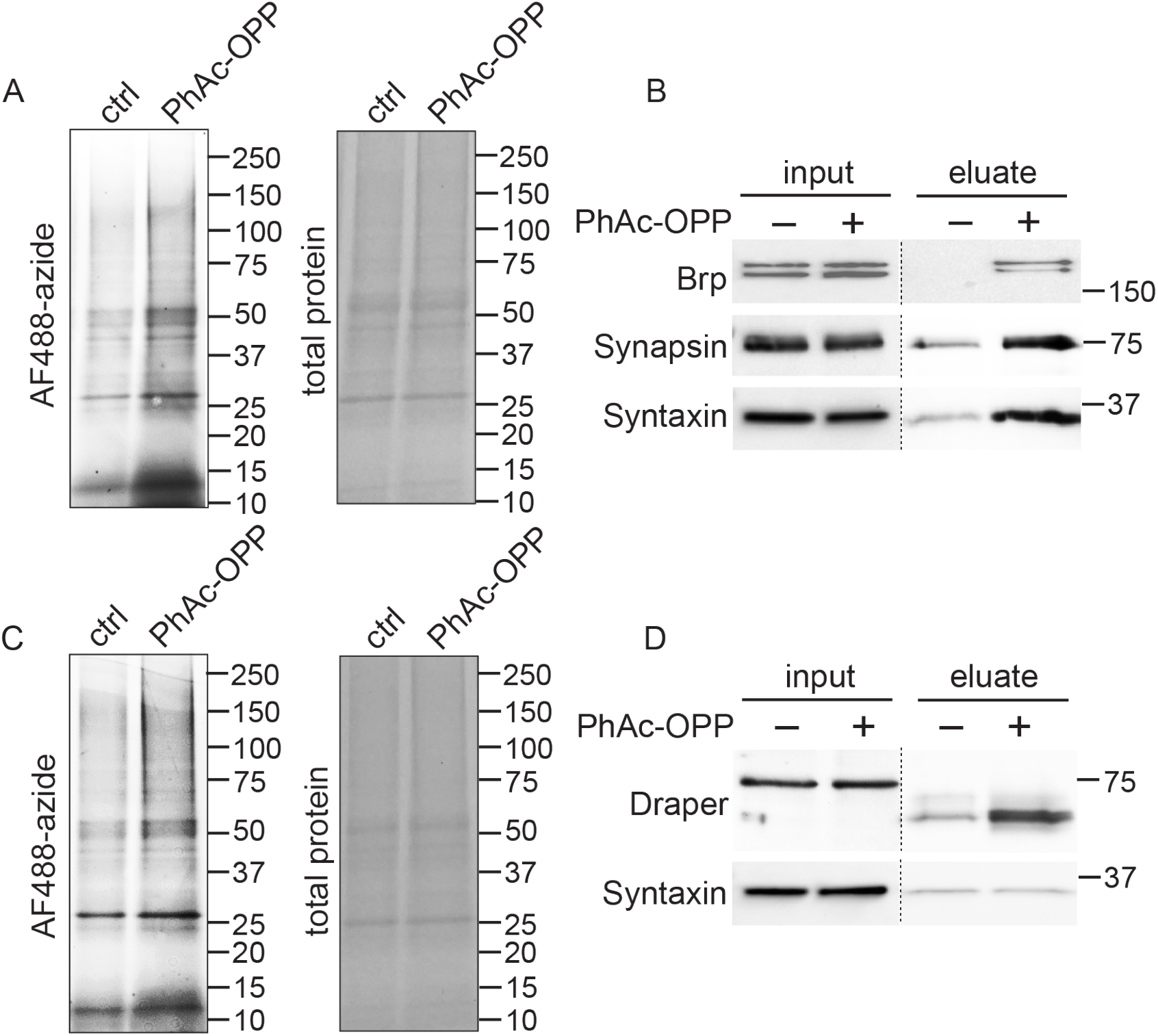
Capture and identification of newly-synthesized proteins from neurons and glia. A, in-gel AF488 fluorescence across a broad range of proteins in fly brain extracts expressing PGA pan-neuronally and following incubation with PhAc-OPP but not vehicle control. B, enrichment of neuron-specific proteins following PhAc-OPP incubation, conjugation of OPP-labeled protein to desthiobiotin azide and neutravidin pull-down. C, pan-glial PGA expression (*Repo-GAL4/UAS-PGA*) promotes broad OPP protein labeling in brain extracts following PhAc-OPP incubation but not vehicle control. D, enrichment of the glial-specific protein Draper (with apparent mol. weight shift between input and eluate) from *Repo-GAL4/UAS-PGA* extracts, but not the neuron-specific protein Syntaxin. Data are representative of three independent experiments.

### No effect of cell type-specific PGA expression on adult fitness or survival

Having found that PGA expression coupled to PhAc-OPP incubation can be used to visualize and capture the nascent proteome in a cell type-specific manner, we sought to determine whether PGA expression in flies affects their development, function or survival. Toward this goal, we crossed *UAS-PGA* flies to pan-neuronal, pan-glial or ubiquitous *GAL4* driver strains and assessed the resulting progeny. Startle-induced negative geotaxis behavior is dependent on a functionally-intact nervous system and declines progressively with age in flies (24, 25). Neither pan-neuronal nor pan-glial PGA expression affects negative geotaxis performance across age (Figure 4A), while ubiquitous PGA expression causes a pronounced deficit in 3-week-old flies but not at a more advanced age in 6-week-old flies (Figure 4B). In accordance with this, ubiquitous PGA expression impairs adult survival, resulting in a ∼20% decrease in median lifespan (Figure 4D and Table 1), whereas no lifespan shortening is seen following pan-neuronal or pan-glial PGA expression (Figure 4C and Table 2). In fact, lifespan appears to be slightly extended by pan-glial PGA (Figure 4C and Table 2). Beside its negative effect on function and survival, ubiquitous PGA expression is also associated with major larval lethality suggesting that development is perturbed, while larval and pupal survival are unaffected by pan-neuronal or pan-glial PGA expression (Figure 4-figure supplement 1) and eclosion occurs at the expected Mendelian frequency. Collectively, these data suggest that PGA expression within the brain is well tolerated across the fly lifespan, and raise the possibility that PGA expression in an unknown tissue outside of the nervous system has a negative impact on development, function and adult survival.

**Table 1.** Ubiquitous PGA expression effect on survival.

| **Genotype** | **N** | **Median lifespan (d)** | **Mean lifespan (d)** | **Log-rank (vs PGA/+ ctrl)** |
| --- | --- | --- | --- | --- |
| *UAS-PGA/+* | 142 | 54 | 53.1 | - |
| *Actin5C-GAL4/UAS-PGA* | 110 | 44 | 45.6 | .006 |

**Table 2.**
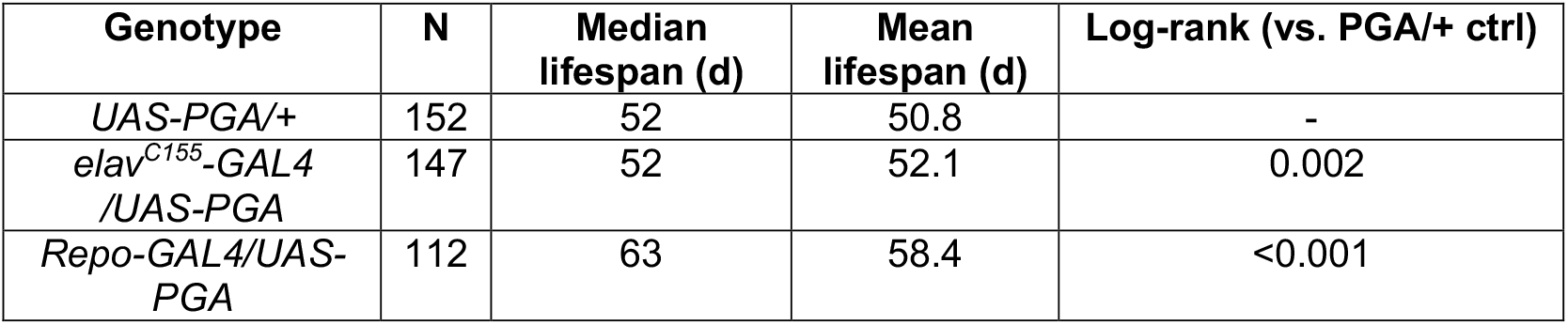
Neuronal and glia PGA expression effect on survival.

| **Genotype** | **N** | **Median lifespan (d)** | **Mean lifespan (d)** | **Log-rank (vs. PGA/+ ctrl)** |
| --- | --- | --- | --- | --- |
| *UAS-PGA/+* | 152 | 52 | 50.8 | - |
| *elav<sup>C155</sup>-GAL4/UAS-PGA* | 147 | 52 | 52.1 | 0.002 |
| *Repo-GAL4/UAS-PGA* | 112 | 63 | 58.4 | <0.001 |

**Figure 4.**
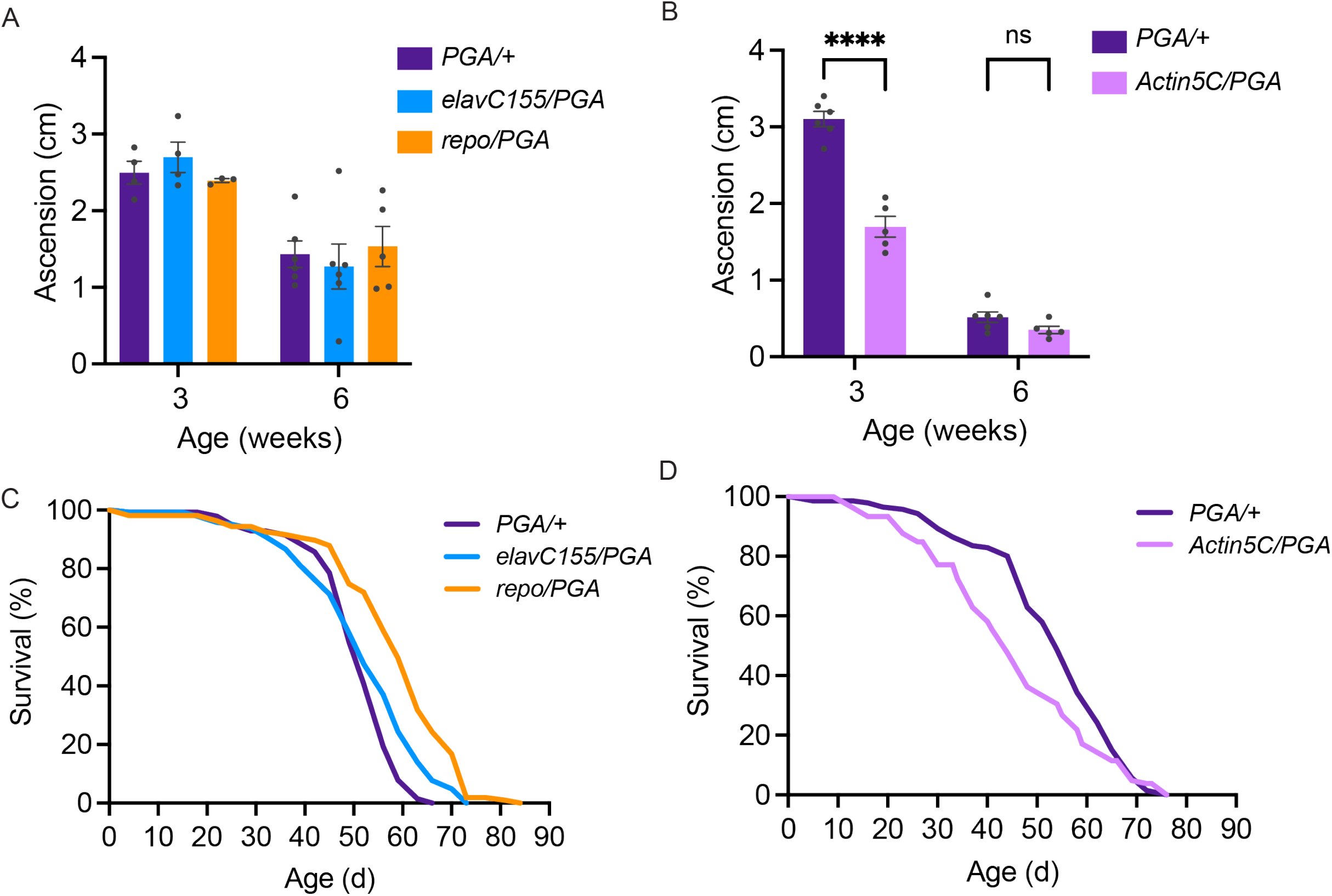
No deleterious effect of cell-specific PGA expression. A, no significant effect of pan-neuronal or pan-glial PGA expression on negative geotaxis behavior at 3 and 6 weeks of age (two-way ANOVA for effect of age (p<0.0001) and genotype, (n.s.), n = 3-6 groups of 25 flies/genotype/age). B, Significant effect of ubiquitous PGA expression on negative geotaxis behavior at 3 weeks of age (two-way ANOVA for effect of age (p<0.0001) and genotype (p<0.0001), Bonferroni post-test, **** p<0.0001, n = 5-6 groups of 25 flies/genotype/age). C, survival is slightly extended upon pan-neuronal or pan-glial PGA expression. D, significant effect of ubiquitous PGA expression on survival. For C and D, see Tables 1 and 2 for experimental n, lifespan metrics and log-rank (Mantel-Cox) comparison results. Data are mean ± SEM. See also Figure 4-figure supplement 1.

### Brain explant labeling can capture *in vivo* protein synthesis states

In order to measure cell type-specific protein synthesis under various physiological or pathological conditions, it is important to know whether POPPi can be used to capture *in vivo* protein synthesis states in newly-isolated brains. To address this question, we examined neuronal protein synthesis labeling in the brains of young and aged flies. It is well established that a widespread age-related decline in protein synthesis occurs in the tissues of numerous organisms including *Drosophila*, when measured across the whole body (26) or in heads (27), and that the decline is due to reduced mRNA translation as well as transcript abundance. To determine whether this same decline is seen specifically in the brains of aging flies, we measured global protein synthesis in young (4-day-old) and aged (21-day-old) wild type fly brain and observe a substantial decrease in radiolabeled protein in aged flies, indicative of decreased protein synthesis in aged fly brain (Figure 5A). Next, using POPPi in which PGA was expressed pan-neuronally, we were able to observe a decline in neuron-specific bulk protein synthesis in the aging fly brain (Figures 5B and C). Strikingly, neuronal protein synthesis was reduced by almost 50% in 3-week-old *elavC155-GAL4/UAS-PGA* flies relative to young (4-day-old) flies of the same genotype (Figures 5B and C).

**Figure 5.**
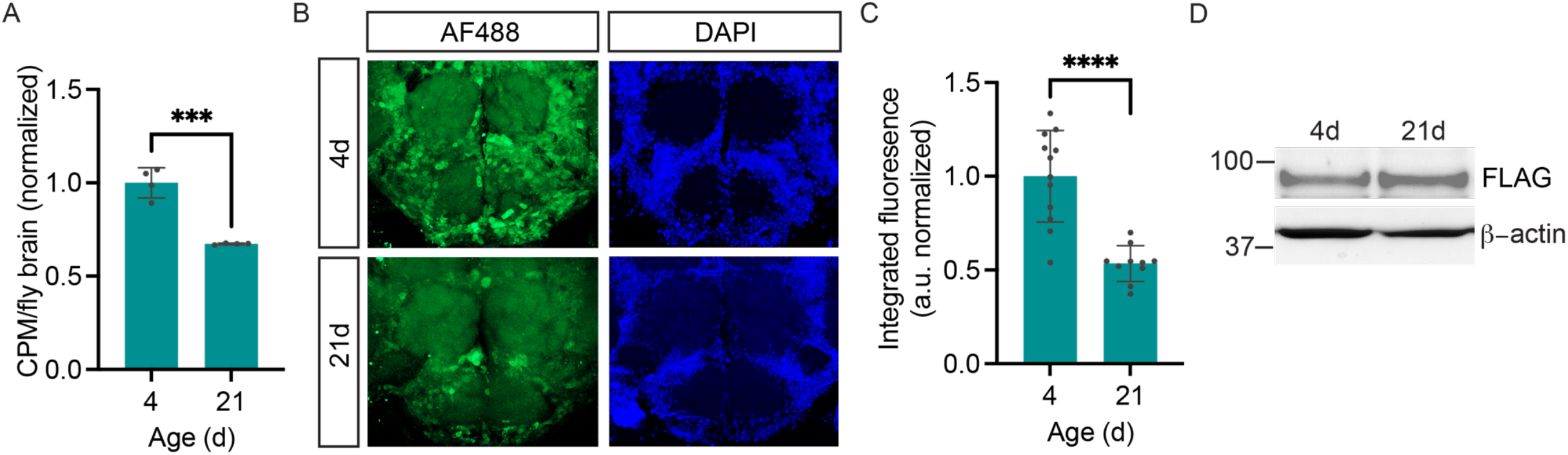
Age-dependent decline in neuronal protein synthesis rate. A, Significant age-related decline in whole brain protein synthesis in aging flies, measured by ^35^S-met/cys labeling (Student’s t-test, ***p<0.001, n =4 groups of 8 brains/age). B, Nascent proteome labeling in young (4-day-old) vs. aged (21-day-old) fly brains expressing pan-neuronal PGA following PhAc-OPP incubation. C, significant effect of aging on protein synthesis (Student’s t-test, p<0.0001, n =10-12 brains/group). Fluorescence signal at each age were derived by subtracting from the mean of a no-label control brain population tested in parallel. D, PGA expression is comparable in 4- and 21-day old *elavC155-GAL4/UAS-PGA* fly brains. Data are mean ± SEM.

Measurement of PGA expression confirmed that this age-related decline in protein labeling was not due to a loss of PGA expression in aged flies (Figure 5D). These data support the conclusion that PGA expression coupled to PhAc-OPP incubation can be used to achieve quantitative assessment of protein synthesis and that an age-dependent decline in protein synthesis is effectively captured by this method.

## DISCUSSION

POPPi is a new method to rapidly visualize and capture cell-specific nascent proteomes in whole *Drosophila* brains. This method extends the demonstrated capability of PGA-dependent OPP labeling in cultured neurons (12) to proteome labeling within complex nervous system tissue. OPP efficiently labels nascent proteomes within the fly brain and this is substantially blocked by the protein synthesis inhibitor cycloheximide (Figures 1C and D), indicating that *de novo* protein synthesis is key for labeling to occur. We show that our strategy is versatile and can be used to visualize nascent protein synthesis across all CNS neurons, specific to the small population of dopaminergic neurons or limited to glia (Figure 2). We also demonstrate that POPPi can be used to capture neuronal or glial proteomes and enrich for proteins of interest within those cell populations (Figure 3). Global protein synthesis is stable for at least 8 hours in isolated whole brain preparations (Figure 1A), supporting an ability to capture *in vivo* protein synthesis states using this approach. Consistent with this, we were able to detect an age-related decline in bulk neuronal protein synthesis in the fly brain using PhAc-OPP (Figure 5). Future efforts will be focused on coupling proteome enrichment to mass spectrometry to interrogate the effects of physiological or pathological stimuli on the proteome at the level of individual proteins.

Spatially-resolved proteomic studies have lagged behind transcriptomic studies in part because issues with limited RNA starting material can be overcome by PCR amplification of cDNA, while no equivalent exists for protein. Nonetheless, flies can be rapidly bred to vast numbers and we anticipate that the facile scalability of *Drosophila* can be leveraged for proteomic studies that focus on small cell populations. PGA transgenic flies are well-suited for CNS proteomic studies because PGA is robustly expressed in adult, larval and pupal brains (Figure 1-figure supplement 1) as well as via neuronal and glial GAL4 drivers (Figure 2-figure supplement 1). Accordingly, while our efforts centered on characterizing proteome labeling within the adult CNS in this study, we anticipate that our chemical genetic approach should be applicable to examining CNS proteomes during fly development. *Drosophila* have been widely used in genetic studies of nervous system development (28), function (29), aging (30) and disease (31). Major insights from these studies highlight strong conservation with mammalian biology and have spurred diverse proteomic analyses of gene expression in the fly nervous system (21, 32-34) and interest in using flies to model the role of translational regulation in nervous system disease (35-37). In addition to the work on diseases such as fragile X syndrome (35), frontotemporal dementia (37) and diseases associated with aminoacyl-tRNA synthetase mutations (36) by others, we recently showed how aberrant translation contributes to neurodegenerative phenotypes caused by the common Parkinson’s disease-causing mutation LRRK2 G2019S in *Drosophila* and iPSC-derived dopamine neurons (38-40), adding to emerging evidence of translational dysregulation in models of Parkinson’s disease (41). Hence, we foresee many opportunities for PGA-expressing flies to generate insight into nervous system function and disease through cell type-specific proteomic studies.

One potential concern surrounding the use of puromycin labeling is that its incorporation into NPC causes chain termination, therefore at high-enough doses it could impede cellular protein synthesis (13). This may be circumvented by finding an optimal concentration of puromycin (or analog) which permits sufficient labeling for detection while having minimal impact on total protein synthesis. We assessed this in *Drosophila* brains and found that 100 μM PhAc-OPP allowed us to obtain robust and rapid proteome labeling while not observing significant deficits in global protein synthesis that were seen at a higher concentration (Figure 2E). We assessed whether ectopic PGA expression in fly cell populations perturbs organism development, function or survival. In contrast to ubiquitous PGA expression, neither neuronal nor glial PGA expression has any discernible negative impact on development, adult function or survival (Figure 4 and Figure 4-figure supplement 1). Hence, when expressed in individual nervous system cell populations, PGA expression seems to be well tolerated. This may provide an advantage over existing NCAA-based cell-specific labeling strategies, where chronic ANL feeding to flies prior to proteomic assessment significantly impairs fly eclosion as well as negative geotaxis behavior in adults (10).

There are additional concerns over how chronic NCAA feeding in this approach might affect protein abundance and overall proteome makeup. For example, replacement of methionine with the NCAA L-azidohomoalanine (AHA) was seen to cause substantial changes in the abundance of numerous proteins in HeLa cells while AHA incorporation into the developing mouse proteome results in altered expression of about 10% of proteins (42, 43). It is also currently unclear whether long-term NCAA administration disrupts eukaryotic metabolism given that AHA and L-homopropargylglycine (HPG) were found to alter global metabolism in E. coli (44). Taken together with the fact that NCAA labeling requires extended dietary methionine depletion, lengthy feeding with amino acid analogs (9, 10) and that NCAA are not incorporated equally across the proteome, there are significant caveats associated with this labeling strategy that can be avoided using POPPi.

While here we focused on protein synthesis labeling in intact isolated brains, this method should in theory be amenable to proteome labeling in tissues throughout the body, wherever PGA can be adequately expressed. As many tissues cannot be isolated and effectively sustained, this may also require systemic administration of PhAc-OPP via dietary feeding. The success of tissue-specific labeling beyond the nervous system may therefore hinge on whether PhAc-OPP can be administered dietarily and at amounts that yield sufficient OPP conversion and nascent protein incorporation in cells of interest, which is yet to be determined. Despite this uncertainty, the ability to rapidly quantify protein synthesis in cell populations within the CNS established here opens up many possibilities to address important questions pertaining to nervous system development, function, aging and disease.

In summary, we provide a new labeling method for rapidly visualizing, capturing and quantifying cell type-specific nascent proteomes within the *Drosophila* brain. We believe this method will be a powerful tool for studying the role of the proteome and translational control in nervous system function with cellular resolution.

## MATERIALS AND METHODS

### Drosophila stocks and culture

PGA-expressing flies were generated by subcloning N-terminal FLAG tagged full-length PGA cDNA (gift of C. Doe) into the fly transformation vector pUAST between KpnI and EcoRI restriction sites. After sequence verification and successful construct expression following transient transfection of *UAS-PGA* and *GAL4* into *Drosophila* S2 cells, the construct was microinjected into w^1118^ fly embryos (Bestgene, Inc). Transgenic FLAG-PGA expression was confirmed by FLAG Western blotting of adult head extracts after crossing all generated *UAS-PGA* lines to *Actin5C-GAL4* (ubiquitous), *elavC155-GAL4* (pan-neuronal) and *repo-GAL4* (pan-glial). The following strongest-expressing lines were used throughout the study: *UAS-PGA-1* for ubiquitous expression and *UAS-PGA-3* for neuronal or glial expression. The *repo-GAL4, UAS-mCD8::mCherry* recombinant line was a gift from M. Freeman and all other lines were obtained from the Bloomington *Drosophila* Stock Center: *TH-GAL4* (line 8848); *elav*^*C155*^*-GAL4* (line 458); *repo-GAL4* (line 7415); *Act5c-GAL4* (line 25374). All flies were reared and aged at 25°C/60% relative humidity under a 12 h light-dark cycle on standard food medium.

### PhAc-OPP synthesis and preparation

PhAc-OPP was synthesized from chemical precursors, analyzed by proton NMR, analytical thin layer chromatography and purified by flash chromatography as previously described (12).

To prepare PhAc-OPP for use, a stock solution (20 mM dissolved in DMSO) was diluted to 100 μM in Schneider’s *Drosophila* medium (with L-glutamine and sodium bicarbonate), bath sonicated for 6 min, vortexed continuously for 30 sec prior to adding the proteasomal inhibitor MG132 (60 μM) and then kept at RT until use.

### Adult brain preparations and ^35^S-methionine/cysteine assessment of de novo protein synthesis

Adult brains were isolated and maintained for protein synthesis assessment in Schneider’s *Drosophila* medium (formulated for *Drosophila* cells and tissues and supplemented with L-glutamine and sodium bicarbonate), based on previously reported methods (15, 18). Prior long-term culture of *Drosophila* CNS explants has often included the addition of insulin and/or serum to the culture medium (15, 18). As global protein synthesis was maintained at a constant rate for 8 hours of *ex vivo* culture in the absence of exogenously added insulin and serum (Fig 1A) and as insulin/serum might artificially stimulate protein synthesis, we omitted them for our short-term rapid protein labeling approach. Adult fly brains (8-10 per genotype) were harvested in Schneider’s medium and transferred to Schneider’s medium containing ^35^S-methionine/cysteine (2 mCi/ml) to metabolically label newly-synthesized protein. Brains were incubated for 30 min at 25°C with gentle orbital shaking, washed twice in 1 ml of PBS then flash frozen. Brains were homogenized in modified RIPA extraction buffer (50 mM Tris-HCl pH 7.4, 150 mM NaCl, 100 mM EGTA, 1% NP-40, 0.1% SDS, protease inhibitor cocktail) on ice using a pestle gun. After a 30 min incubation on ice, lysates were centrifuged at 14,000 x g for 15 min and the supernatant was retained. Protein was precipitated by the addition of methanol and heparin (lysate:heparin (100mg/ml):methanol volume ratio of 150:1.5:600), centrifuged at 14,000 x g for 2 min, the supernatant was removed, and the pellet was air dried. The protein pellet was resuspended in 8 M urea/150 mM Tris, pH 8.5, and ^35^S-methionine/cysteine incorporation was measured by liquid scintillation counting (counts per minute, CPM) and normalized to the number of brains.

### Effect of PhAc-OPP on global protein synthesis

Adult fly brains (8-10 per genotype) from the standard laboratory control strain *w*^*1118*^ were harvested in Schneider’s medium and transferred to Schneider’s medium containing ^35^S-methionine/cysteine (2 mCi/ml) and PhAc-OPP at the indicated concentrations. Brains were incubated for 2h at 25°C with gentle orbital shaking, washed twice in 1 ml of PBS then flash frozen. Thawed brains were homogenized and processed for protein precipitation and measurement of ^35^S incorporation by scintillation counting as described above.

### Rapid OPP/PhAc-OPP labeling of newly-synthesized CNS proteins

To enable rapid labeling of nascent CNS proteins, adult *Drosophila* brain explants maintained in Schneider’s *Drosophila* medium were incubated with OPP or PhAc-OPP. Brains (∼15 for immunocytochemistry, 100 for enrichment and immunoblotting) were harvested in Schneider’s medium and then immediately transferred to Schneider’s medium containing OPP (50 μM unless otherwise indicated) or PhAc-OPP (100 μM unless otherwise indicated) and MG-132 (60 μM) for 2h (unless otherwise indicated) at 25°C with gentle orbital shaking. Brains were washed twice briefly in 1 ml PBS and then immediately processed for immunocytochemistry or enrichment and immunoblot detection as described below.

### Immunocytochemical detection of OP-puromycylated protein

OP-puromycylated brains were fixed for 20 min in 4% paraformaldehyde in PBS-T (PBS, pH 7.4 containing 0.3% Triton-X-100) at RT with gentle rocking, washed once in chilled PBS-T (5 min) and once in chilled PBS (5 min). AF488 picolyl azide conjugation to OPP was carried out via CuAAC click reaction. A click working reagent was prepared containing TBTA (200 μM), freshly prepared Cu(I)Br (0.5 mg/ml), AF488-picolyl-azide (0.1X concentration) and 1X concentration OPP reaction buffer (the last two reagents derived from the Click-iT™ Plus OPP Alexa Fluor™ 488 Protein Synthesis Assay Kit (Invitrogen) and at concentrations relative to those recommended in the manufacturer’s instructions). The click working reagent was vortexed for 30 sec then applied to brains and incubated for 30 min at 25°C with gentle orbital shaking and protection from light.

Brains were washed briefly with 1 ml of Click-iT® Reaction Rinse Buffer and either whole-mounted or additionally immunostained for cell markers or DAPI stained prior to mounting. Brains were imaged on a Zeiss LSM900 confocal microscope. For in-gel fluorescence assays, OPP/PhAc-OPP treated brains were lightly fixed (4% PFA/10 min), click-conjugated to AF488 picolyl azide as described above then lysed in de-crosslinking buffer (300 mM Tris-HCl/2% SDS/protease inhibitors) for 2h at 60 °C prior to SDS-PAGE, gel fixation (40% methanol/10% acetic acid), 30-minute washing in ddH_2_O, AF488 detection and subsequently total protein visualization using colloidal blue stain.

### Immunoblot detection of enriched OP-puromycylated protein

The protocol is modified from Marter et al. (45). Brains were thawed on ice and homogenized in 100 μL of homogenization buffer (0.5%SDS/PBS containing 2X Complete EDTA-free Protease Inhibitor Cocktail). Samples were incubated on ice for 20 min with mixing every 5 min, at 95 °C for 5 min, then on ice for 5 min. Triton-X-100 was added to a final concentration of 0.2% and pre-equilibrated neutravidin agarose (25 μL/sample) was added to pre-clear lysates of endogenous biotinylated proteins via sample incubation on a nutator at 4 °C for 1h. Samples were centrifuged (3000 x g, 5 min, 4 °C) and to the supernatant, click chemistry reagents were added as follows: TBTA (final concentration 200 μM), desthiobiotin azide (20 μM), freshly prepared Cu(I)Br (0.5 mg/ml). On each addition, samples were vortexed for 10 sec. Samples were incubated overnight on a nutator at 4°C, centrifuged at 3000xg for 5 min/4 °C then the supernatant was diluted in PBS with 2X protease inhibitor cocktail to a final volume of 300 μl and desalted (Zeba spin desalting columns). Total protein concentration in lysates was determined by BCA assay to standardize input for neutravidin agarose enrichments. Neutravidin agarose was equilibrated in 1% NP-40/PBS for three washes and 1% NP-40 was added to desalted lysates for 20 min on ice. After removing a portion of the lysate for input analysis, samples were added to equilibrated neutravidin agarose (50 μL/sample) and incubated overnight at 4 °C on a nutator. Samples were centrifuged (3000 x g, 5 min, 4 °C) and washed in 1% NP-40/2X protease inhibitor cocktail/PBS 5 times then in PBS for 3 additional washes, with 2 min of end-over-end mixing at each wash step. Desthiobiotin-conjugated protein was eluted from neutravidin agarose by incubating in 8 mM biotin for 1h in a thermomixer at 25 °C/1200 rpm shaking. Laemmli buffer was then added to eluate and input for downstream analysis by immunoblotting.

### Adult survival

Adult females (0–3 days of age, 100–150 flies per genotype, selecting against newly-eclosed flies) were collected under brief anesthesia and transferred to fresh food vials at 25 flies per vial. Flies were then transferred to fresh food vials every 3–4 days throughout the experiment and dead or censored (escaped or stuck in food) flies were counted during each transfer to fresh food.

### Negative geotaxis behavior

Cohorts of 75–100 female flies (0–3 days old, selecting against flies with visible signs of recent eclosion) were collected under brief anesthesia and transferred to fresh food vials to recover (25 flies/vial). Flies were aged for six weeks with transfer to fresh food twice per week. On the day of testing, flies were transferred to empty vials, allowed 1 min to rest and then tapped to the bottom of the vial three times within a 1 sec interval to initiate climbing. The position of each fly was captured in a digital image 4 sec after climbing initiation. Automated image analysis was performed using the particle analysis tool on Scion Image to derive x–y coordinates for each fly thus providing the height climbed, as previously described (24). The performance of flies in a single vial was calculated from the average height climbed by all flies in that vial to generate a single datum (N = 1). Performance of each line was then derived from the average scores of 5-6 vials tested for the line (N = 5-6).

### Western blotting

Samples were electrophoresed on 4-20% Tris-Glycine gradient gels and transferred to nitrocellulose membrane for immunoblotting using the following antibodies:

From the Developmental Studies Hybridoma Bank: Brp (nc82) 1:50; Synapsin-1 (3C11) 1:500; Syntaxin (8C3) 1:500; Drpr (1:1 mix of 5D14:8A1) 1:400; Repo (8D12) 1:200. Other antibodies used were FLAG (M2) (Sigma) 1:500; TH (Immunostar) 1:1000; Actin-HRP (Sigma) 1:1000; Ubiquitin (P4D1) (Cell Signaling Technology) 1:1000; GAPDH (GA1R) (ThermoFisher Scientific) 1:10,000.

### Statistical analysis

Quantified data are mean ± SEM, and individual data points are plotted for data with *n* <10. Sample sizes for time-course experiments were determined based on evidence from pilot experiments. Statistical analysis details for each individual experiment are described in figure legends, including the number of flies or number of groups of flies used (*n*), statistical tests (unpaired two-tailed Student’s *t* test, ANOVA or two-way ANOVA) and Bonferroni post hoc analysis with associated *p* values. All statistical analyses were performed using GraphPad Prism except Log-rank (Mantel-Cox) tests for lifespan comparisons performed in SPSS.

## Data availability

All data generated or analyzed during this study are included in the manuscript main Figures or Figure supplements. Source data are provided for Western blots and gels.

## Materials availability

Newly created *Drosophila* lines will be shared upon request and upon final manuscript publication.

## ACKNOWLEDGMENTS AND FUNDING SOURCES

We thank Richard Goodman for initial discussions on the project, Chris Doe and Sen-Lin Lai for providing PGA cDNA, the OHSU Advanced Light Microscopy Core for microscope use and the OHSU Medicinal chemistry core for PhAc-OPP synthesis. The following antibodies were obtained from the Developmental Studies Hybridoma Bank, created by the NICHD of the NIH and maintained at The University of Iowa: Brp nc82 and Synapsin 3C11 (developed by E. Buchner), Syntaxin 8C3 (S. Benzer and N. Colley), Draper 8A1 and 5D14 (M. Logan) and Elav 9F8A9 (G. Rubin). This work was funded by OHSU Neurology Foundation Funds (I.M.).

## FIGURES LEGENDS

**Figure 1-source data file**. Source gel images for Fig. 1E

**Figure 1-figure supplement 1 -source data file**. Source western blots for Figure 1-figure supplement 1C.

**Figure 2-source data file**. Source western blots for Figure 2F.

**Figure 3-source data file**. Source western blots for Figure 3.

**Figure 5-source data file**. Source western blots for Figure 5D.

## FIGURE SUPPLEMENTS AND TABLES

**Figure 1-figure supplement 1.**
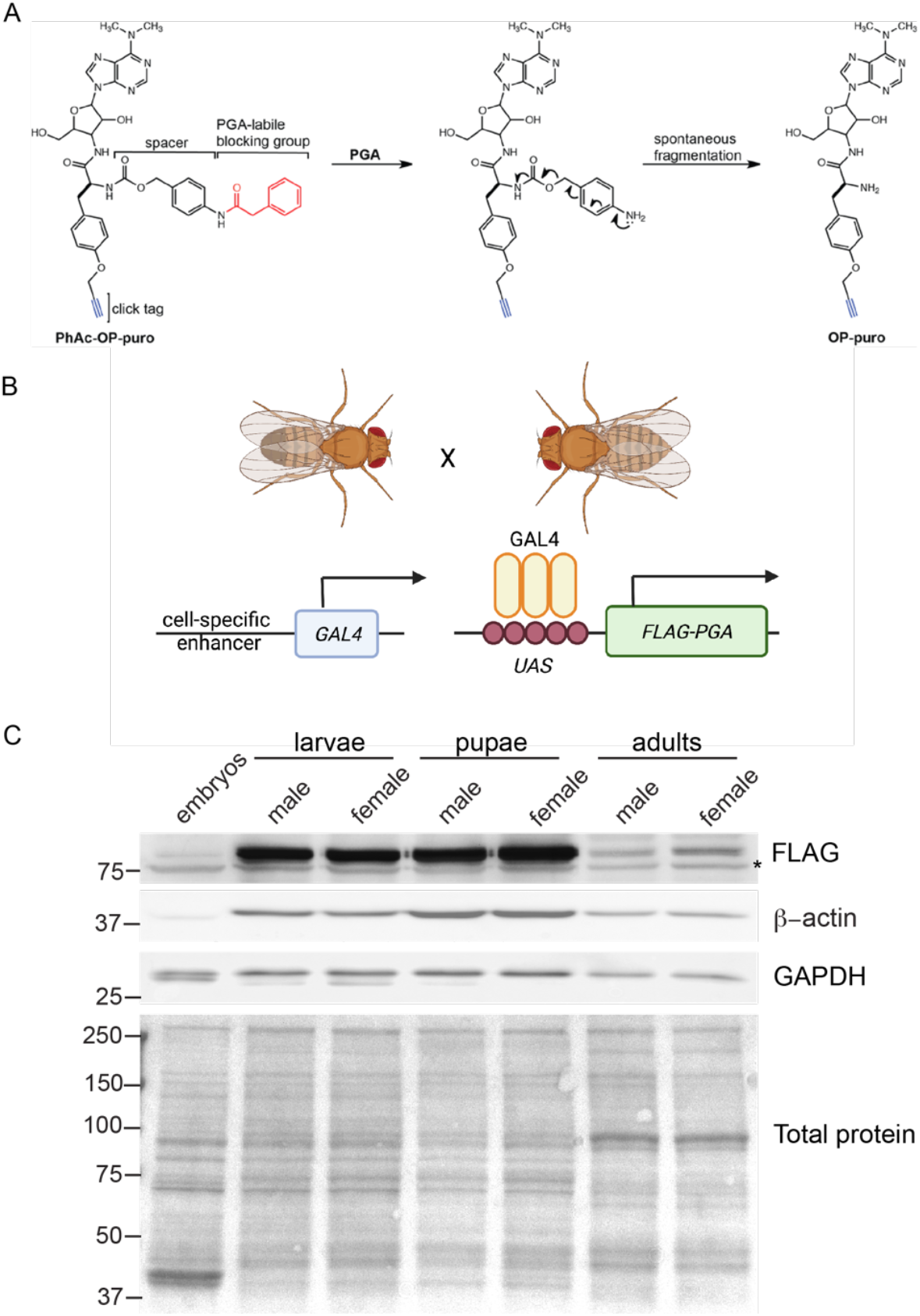
PGA-dependent conversion of PhAc-OPP to OPP in *Drosophila*. A, scheme illustrating the conversion of PhAc-OPP to OPP by penicillin G acylase (PGA). PGA catalyzes removal of the phenylacetyl blocking group and the carbamate spacer then undergoes spontaneous fragmentation to generate OPP. Reprinted (adapted) with permission from R.M. Barrett et al., ACS Chemical Biology 2016 (12). Copyright 2016 American Chemical Society. B, scheme illustrating expression of N-terminally FLAG-tagged PGA under a cell-specific driver of choice using the binary GAL4/UAS system. C, PGA expression levels within whole embryos or the brains of L3 larvae, pupae or adult flies expressing PGA via the ubiquitous *Actin5C-GAL4* driver. Asterisk, non-specific band. Both loading controls (β-actin and GAPDH) are divergent in expression, size and/or number of bands between embryos, larvae, pupae and adults, hence a Ponceau stain is shown for protein loading. Illustration in B created on Biorender.

**Figure 1-figure supplement 2.**
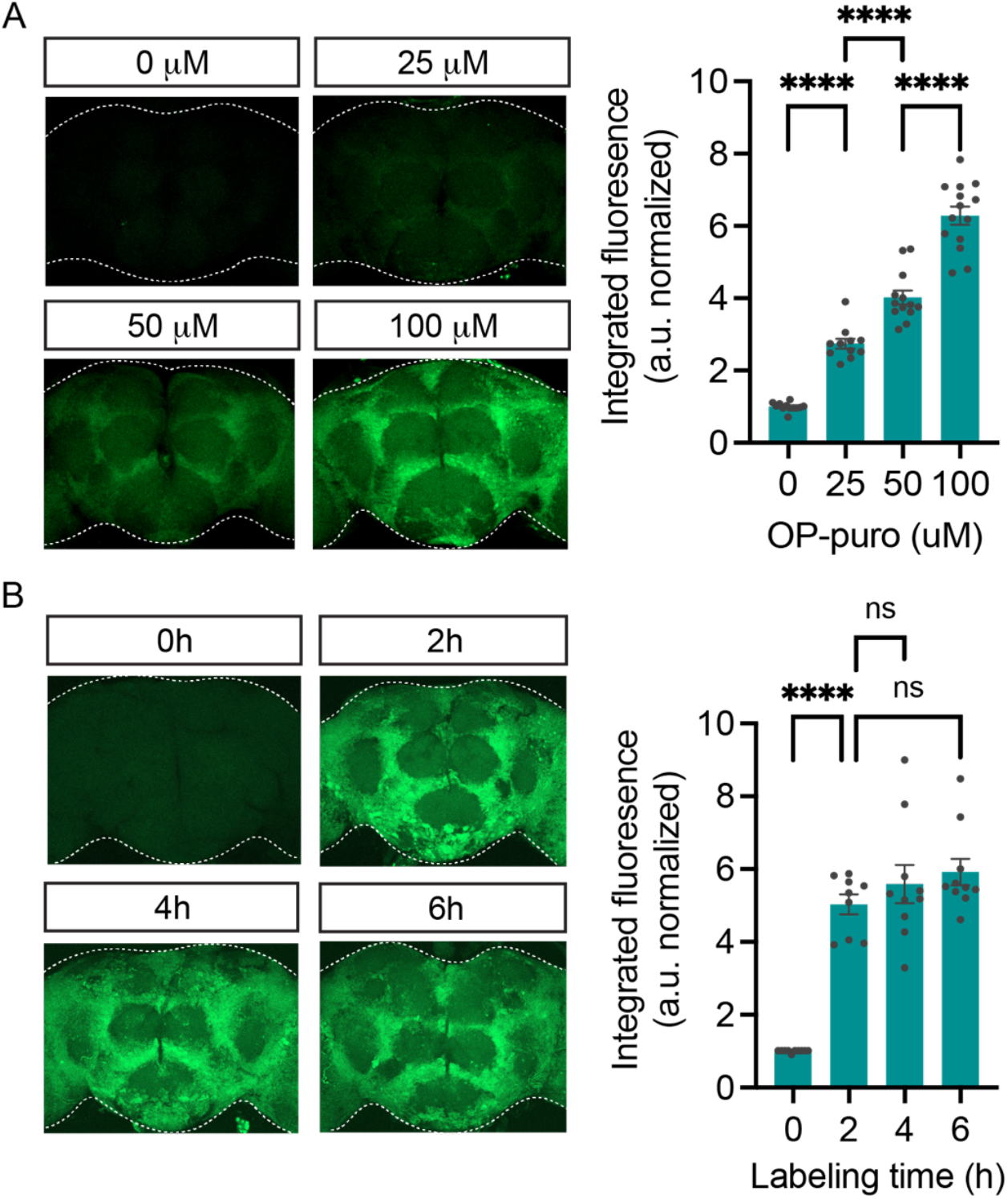
OPP nascent proteome labeling is concentration and time-dependent. A, significant effect of OPP concentration on OPP labeling in standard laboratory control flies (*w*^*1118*^, ANOVA, p<0.0001, Bonferroni post-tests, **** p<0.0001, n =11-14 brains/group). Labeling time was 2h. B, significant effect of labeling time on OPP labeling in w^1118^ flies (ANOVA, Bonferroni post-tests, **** p<0.0001, n =9-10 brains/group). 50 μM OPP was used. Data are mean ± SEM.

**Figure 2-figure supplement 1.**
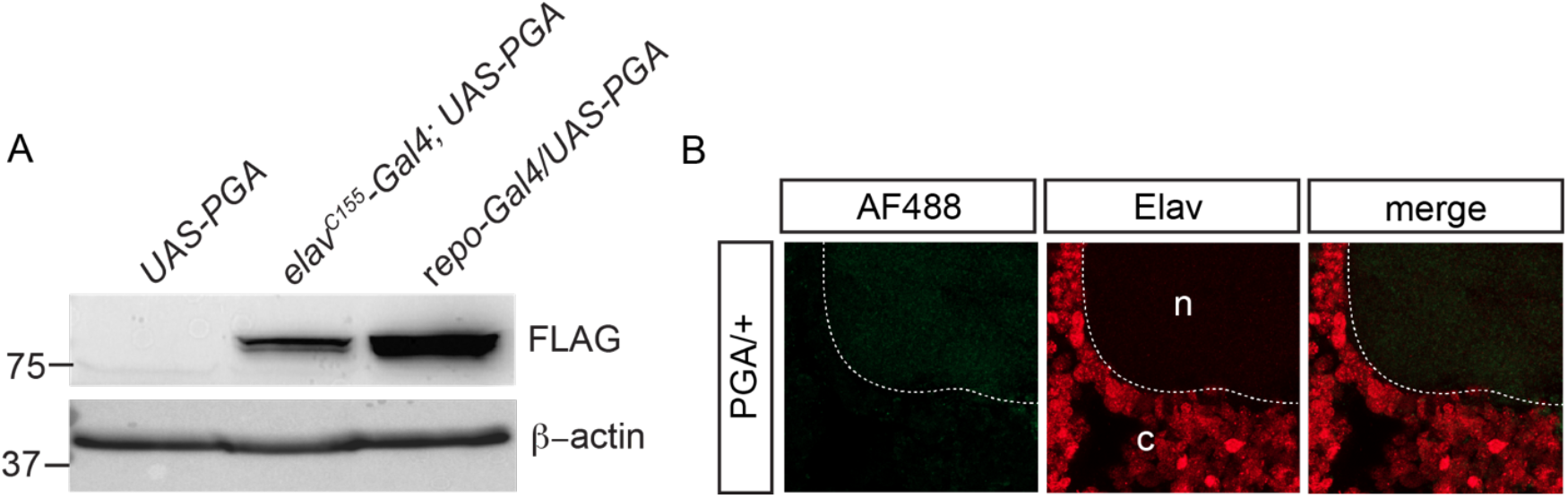
Cell-specific labeling via PGA expression. A, FLAG-PGA expression via *elav*^*C155*^*-GAL4* or *repo-GAL4* in whole brain extracts. B, no PhAC-OPP labeling in *UAS-PGA/+* control fly brains lacking the *elav*^*C155*^*-GAL4* driver, incubated with PhAc-OPP. c, cell cortex showing positive immunostaining for the neuronal nuclear protein Elav. n, neuropil.

**Figure 4-figure supplement 1.**
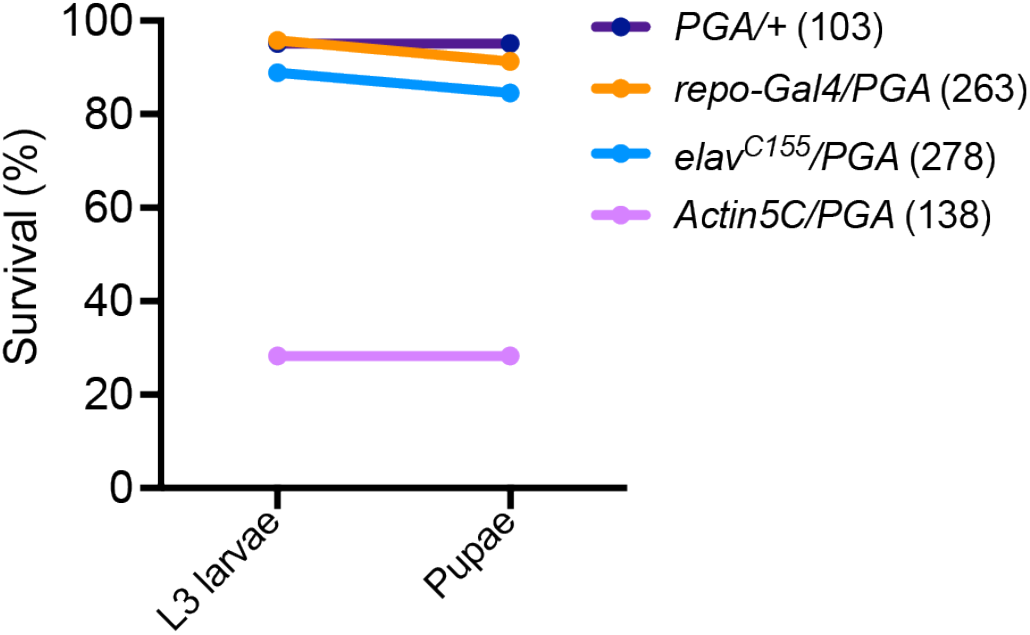
Larval lethality following ubiquitous PGA expression. Numbers in parentheses are number of flies tested.

## Notes

### Competing Interest Statement

The authors have declared no competing interest.

